# Hyperspectral Sensing for High-Throughput Chloride Detection in Grapevines

**DOI:** 10.1101/2024.06.20.599906

**Authors:** Sadikshya Sharma, Christopher Wong, Krishna Bhattarai, Yaniv Lupo, Troy Magney, Luis Diaz-Garcia

## Abstract

Soil salinity affects major viticultural areas worldwide with chloride ions being the primary source of salt toxicity in grapevines. This toxicity impacts vine health and reduces fruit yield and quality. Current breeding efforts to improve grapevine salinity tolerance are limited by the low throughput of available phenotyping methods, which are time-consuming, labor-intensive, and destructive. This study demonstrated that hyperspectral proximal sensing can be utilized as a high-throughput, non-destructive screening technique to identify salinity-tolerant grapevine germplasm. The predictive abilities of two different hyperspectral devices, which varied in price, resolution, and sensitivity, were compared across 23 *Vitis* accessions spanning eight species. Prediction models were built using hyperspectral reflectance and leaf chloride content measured with a lab chloridometer. Three distinct approaches were studied: 1) analyzing the correlation between individual wavelengths and chloride content; 2) employing machine learning models, including Partial Least Squares Regression (PLSR), Random Forest (RF), and Support Vector Machine (SVM), utilizing all wavelengths; and 3) classification-based prediction using Partial Least Squares Discriminant Analysis (PLSDA). Multiple regions in the spectrum, including 613-660 nm, 689-696 nm, and 1357-1358 nm, showed a medium correlation (0.30-0.50) with chloride content in the leaves. PLSR was the most effective machine learning approach, demonstrating moderate predictive capability for chloride content (maximum *R²* = 0.67), though performance varied between the two devices tested. With PLSDA, predictions increased considerably, up to an accuracy of 0.97, depending on the instrument used and the spectral data transformation. Overall, the more expensive and sensitive device with a wider spectral range outperformed the more affordable, shorter-range device. However, when the prediction model was based on classes (chloride excluders vs. non-excluders) rather than chloride content, the differences in prediction abilities were minimal, with both instruments performing very well. This is promising for identifying breeding materials with chloride exclusion capabilities at low cost and high throughput.

## 1 INTRODUCTION

Increasing soil salinity is a global threat to major viticultural areas as the quality and quantity of irrigation water decreases (Ollat et al., 2016). The impact of salinity on crop production is expected to worsen with climate change, particularly in Mediterranean regions (Olesen, 2011). Salinity hinders plant growth and development by causing water stress and cytotoxic effects from the accumulation of ions like sodium (Na^+^) and chloride (Cl^−^), ultimately affecting yield and quality (Calzone et al., 2021). While chloride is an essential plant element, its accumulation beyond a certain threshold leads to physiological disruptions that reduce net photosynthesis, leaf area, and water uptake (Walker, 1994). Chloride toxicity is a major abiotic stressor in arid and water-limited regions where grapes are grown, raising concerns in the viticulture community (Walker, 1994; Walker et al., 2003; Heinitz et al., 2020). *Vitis vinifera* cultivars and many commercial rootstocks are sensitive to chloride concentrations commonly found in viticultural areas, resulting in reduced productivity, low fruit and wine quality (Walker. 1994; Walker et al., 2003). Fortunately, several wild *Vitis* species have been identified as chloride excluders because of their ability to restrict chloride uptake (Heinitz et al., 2020). These species prevent chloride ions from accumulating to toxic levels in leaves and fruit, maintaining vine health and quality (Ollat et al., 2016; Tregeagle et al., 2010). Current methods for measuring chloride exclusion in grapevines are destructive, time-consuming, and low throughput. Typically, this process involves harvesting, drying, and grinding leaves before determining chloride content with a lab-bench chloridometer (Heintz, 2020). From a breeding perspective, this methodology limits the amount of plant material that can be evaluated. Therefore, developing a high-throughput screening method is essential for breeding programs focused on selecting vines with improved chloride-exclusion capabilities.

Hyperspectral reflectance data from leaf surfaces offers valuable insights into the physiological and biochemical status of plants. (Gates et al., 1965; Knipling, 1970; Wong, 2023). Spectral features from leaves are based on how light affects the vibration of the organic molecular bonds, especially, C-H, N-H, and O-H in visible (VIS; 400-700 nm), near-infrared (NIR; 700-1100 nm), and shortwave-infrared (SWIR; 1100-2500 nm) spectral regions (Wong et al., 2023). Understanding the absorption and reflectance of light across spectral features allows for inferences about vegetation physiology, biochemistry, and structure. The VIS spectrum is particularly sensitive to light absorption by pigments such as chlorophyll, anthocyanins, and carotenoids (Blackburn, 2007). The NIR spectral region responds to leaf structure, revealing variations among genotypes or phenological stages (Asner, 1998). Similarly, the SWIR region is sensitive to leaf structure, water content, and phenolic compounds (Kokaly and Skidmore, 2015). In many species, spectroscopy technologies, including NIR and infrared, have been used to determine nutrient content non-destructively (Niederberger et al., 2015), biomass (Jin et al., 2017), water status (Zhang et al., 2021), and stress response (Huang et al., 2012). Notably, using a NIR benchtop spectrometer with dried, pulverized tissue samples, the chloride content in persimmon trees grown in high-salinity orchards was accurately predicted (*R^2^* = 0.93) through PLSR (de Paz et al., 2016). Specific wavelengths, particularly 1527–1563 nm, 1725 nm, and 1995–2091 nm showed strong correlations with chloride content (*R^2^* > 0.56). Despite its predictive strength, this approach remains labor-intensive and destructive. Conversely, similar research on persimmon trees identified the 390-472 nm and 690-692 nm wavelength ranges as strongly associated with chloride accumulation in leaves using a non-destructive method (Visconti and de Paz, 2019). These findings suggest the potential for using portable sensors to infer chloride content in plant tissues non-destructively.

This study investigated the use of hyperspectral proximal sensing for predicting chloride content in several *Vitis* species with potential for rootstock breeding. Considering the typical constraints on breeding budgets, two different hyperspectral devices—differing in cost, spectral range, and resolution—were employed to collect leaf reflectance data from potted grapevines subjected to salt stress. Several techniques for hyperspectral data processing were tested, along with three approaches for predicting chloride content.

### Core ideas

- Hyperspectral reflectance data can moderately predict chloride content in grapevines.
- Different devices and spectral preprocessing methods produced variations in prediction accuracy.
- PLSR predicts leaf chloride content with higher accuracy than RF and SVM
- Classification-based prediction utilizing PLSDA effectively distinguishes between chloride excluders and non-excluders with higher accuracies.
- Hyperspectral phenotyping can enhance selection intensity, ultimately leading to increased genetic gain

## 2 MATERIALS AND METHODS

### 2.1 Plant material and experimental design

Dormant bud cuttings of wild *Vitis* germplasm and commercial rootstocks with varying chloride exclusion capacities were collected from mother vines planted in Davis, California, in December 2022. The accessions were selected based on previous chloride exclusion screenings (Heinitz et al., 2020). The plant materials included selections of the wild species *V. cinerea*, *V. berlandieri*, *V. girdiana*, *V. acerifolia*, *V. treleasei*, *V. arizonica*, *V. mustangensis*, *V.* X *champinii*, *V*. X *doaniana*, and *V. rupestris*, along with two commercial rootstocks, 140 Ru and Ramsey (Table S1). The cuttings were soaked overnight in 1% bleach water, placed in callusing media, and kept in a dark room at 100% relative humidity and 29 °C for two weeks. Following this, the cuttings were cleaned, the top buds were waxed, and placed in a paper sleeve with a 1:1 ratio of callus media: peat moss. The cuttings were kept for an additional four days in a dark room at 100% relative humidity and 29 °C. Subsequently, plants were transferred to a fog room at 27 °C with bottom heat for two weeks before being transplanted into a two-liter pot with fritted clay.

Two similar experiments were conducted as full factorial designs, with five repetitions per genotype:salt treatment during the spring and early summer of 2023. Salt treatments, initiated two months after transplanting, comprised three NaCl levels (0 mM, 50 mM, and 75 mM of NaCl) and lasted for 21 days. Each pot was watered with one liter of the respective saltwater concentration every morning between 7 and 8 AM. Both trials utilized the same 23 accessions, experimental design, and treatments. The salt treatment in trial 1 started on May 17, 2023, and concluded on June 6, 2023, while trial 2 began on June 28, 2023, and ended on July 18, 2023.

### 2.2 Leaf sampling and spectral measurements

Hyperspectral information was captured using two distinct spectral single-point devices, each differing in wavelength range, resolution, size, operability, and cost. The first device SVC HR-1024i (Spectra Vista Corporation in Poughkeepsie, NY, USA), coupled with a leaf clamp (LC-RP Pro), is a field-portable spectroradiometer that records reflectance spectra across 350 to 2500 nm wavelengths; its resolution varies: 1.5 nm between 350 and 1000 nm, 3.8 nm from 1000 to 1890 nm, and 2.5 nm from 1890 to 2500 nm. Despite its larger size, the HR-1024i remains portable. The second device, the NIR-S-G1 (Innospectra Corporation, Hsinchu, Taiwan), captures reflectance within the 900 to 1700 nm range, with a spectral resolution of 3 to 4 nm. This device is significantly smaller and approximately 20 times less expensive than the SVC HR-1024i. For spectral data collection, the fifth leaf from the base of the plant was selected. Measurements were taken between 10 AM and 12 PM on day 1 and day 21 of the experiment. Leaf reflectance was determined by dividing the leaf reflected radiance by a prior white reference (incoming active light source) scan and the scan time was set to one second. A reference scan was collected every 20 leaves or approximately every 10 minutes, to maintain measurement accuracy.

### 2.3 Reference leaf chloride measurement

After 21 days of salt treatment, all leaves and petioles from each plant were collected and stored in a paper bag. The samples were air-dried in a drying room at 50°C for two weeks before being grounded. After 14 days, the samples were further pulverized using a metal rolling pin until a fine powder was obtained. Subsequently, 0.25 g of the fine powder was mixed with 25 ml of deionized water in screw cap bottles, following the protocol of Heinitz et al., 2020. The bottles were shaken at 150 RPM for 1 hour, and the mixture was filtered through an 11 μm filter to obtain a clear solution without leaf particles. The chloride content of the filtered samples was then measured using a silver ion titration chloridometer (Model 926, Nelson-Jameson Inc., Marshfield, WI, USA) following the manufacturer’s calibration protocol with standards of known chloride concentrations. Each sample was measured three times, and the average chloride content, expressed in µmol·g^−1^ of DW, was used for further analysis.

### 2.4 Spectral data analysis

To amplify the signal-to-noise ratio, rectify non-linear spectral trends, and reduce variability caused by scattering, particle size effects, and path length differences in light, raw spectra were processed with 12 preprocessing methods (also called transformations or treatments) using the R package ‘waves’ version 0.2.4 (Hershberger et al., 2021). These preprocessing methods included: 1) standard normal variate (SNV), 2) first derivative (D1), 3) second derivative (D2), 4) standard normal variate and first derivative (SNV1D), 5) standard normal variate and second derivative (SNV2D), 6) Savitzky-Golay filter (SG), 7) standard normal variate and Savitzky-Golay filter (SNVSG), 8) gap=segment derivative (window size = 11; SGD1), 9) Savitzky-Golay filter first derivative (window size = 5; SG.D1W5), 10) Savitzky-Golay filter and first derivative (window size = 11; SG.D1W11), 11) Savitzky-Golay filter and second derivative (window size = 5; SG.D2W5), and 12) Savitzky-Golay filter and second derivative (window size = 11; SG.D2W11).

Chloride content and processed spectral reflectance were analyzed using three distinct approaches. First, Pearson’s correlation (r) was used to examine the relationship between individual wavelengths across the hyperspectral profile and chloride content. Second, three machine learning techniques leveraging the entire spectral profile—PLSR, RF, SVM—were evaluated using the reflectance data alongside the 12 preprocessing methods mentioned earlier. The effectiveness of these models was assessed through squared Pearson’s correlation between observed and predicted values in the validation set (*R^2^*), the root mean squared error of prediction (*RMSE*), the residual prediction deviation (*RPD*), the ratio of performance to inter-quartile (*RPIQ*), the concordance correlation coefficient (*CCC*), and the bias. These indicators collectively provided a comprehensive evaluation of each model’s predictive accuracy and overall fit.

For PLSR, RF, SVM, and PLSDA, five-fold cross-validation was conducted, dividing the data into a 70 % training set and a 30 % validation set. PLSDA models were assessed with 100 retraining iterations to generate new training and validation sets each time, to maximize accuracy (proportion of true positives and negatives relative to the population size) and specificity (true positives relative to the sum of true positives and false negatives) for each model iteration. Leveraging the two separate trials conducted in this study, five different prediction scenarios involving PLSR, RF, and SVM were compared for spectral data obtained from SVC HR-1024i (Figure 1). Scenarios 1 and 2 involved training and validation within the same trial. In scenario 3, data from both trials were combined for training and validation. For scenarios 1 to 3, 70% of the data was used for training, while 30% was reserved for validation. The validation phase involved comparing predictions to the actual chloride content by calculating the squared Pearson’s correlation (*R²*) between observed and predicted values in the validation set. Scenarios 4 and 5 involved training the models with data from one trial and validation on the other trial. In these scenarios, 100% of the data from one trial was used for training, and validation on 100% of the data from the other trial. For spectral data from NIR-S G1, only scenario 1 was tested, as spectral data was collected exclusively from trial 1.

**Figure 1.**
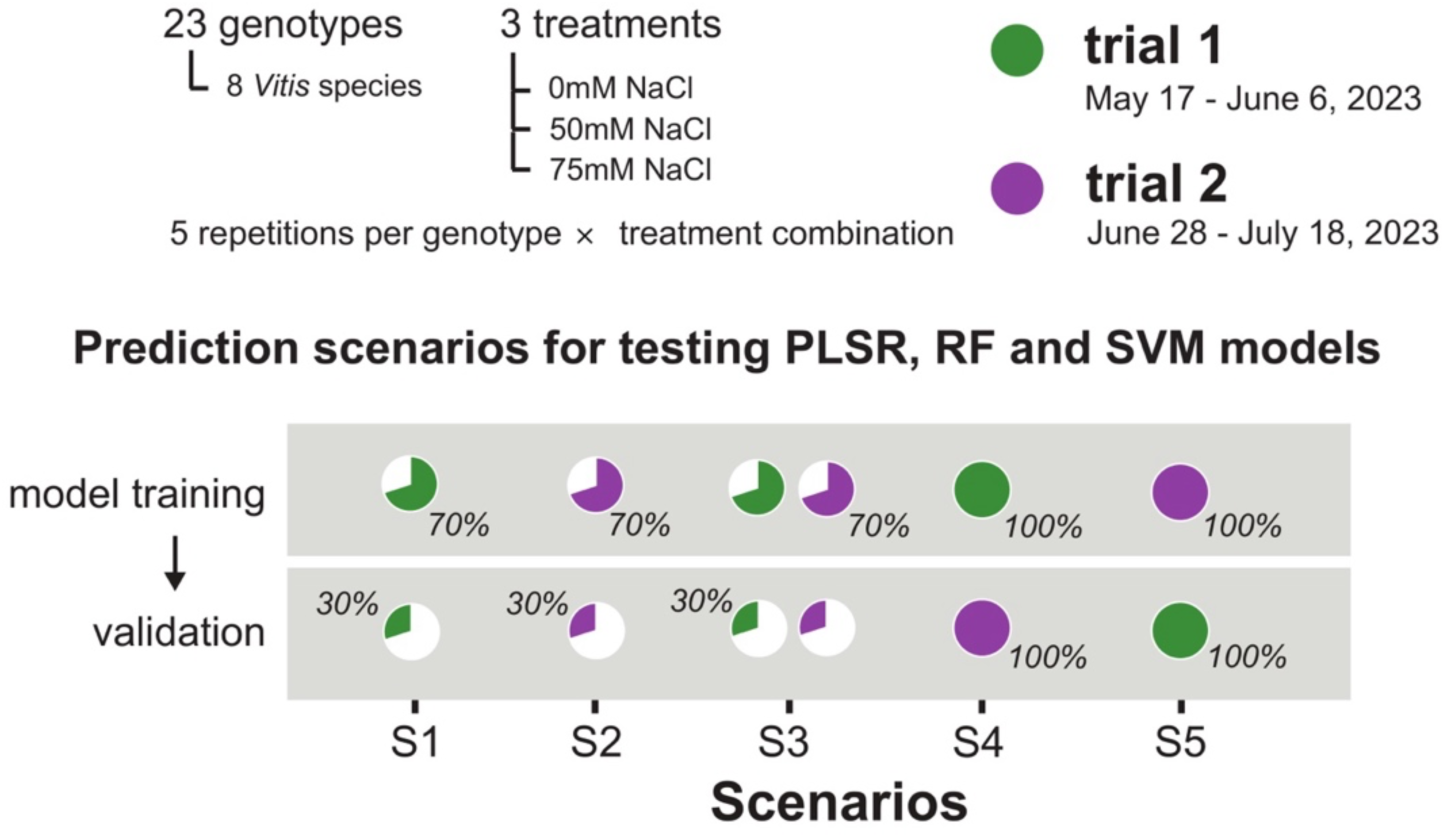
Graphical representation of five prediction scenarios tested using Partial Least Square Regression (PLSR), Random Forest (RF), and Support Vector Machine (SVM)

Furthermore, PLSDA, an adaptation of PLSR for categorical variables, was utilized after discretizing quantitative chloride content values into two groups: chloride excluders and non-excluders. Only scenario 3 was tested using this approach, applied exclusively to plants treated with 50 mM and 75 mM NaCl. Two approaches were compared: The first, the combined treatment approach, considering all treated plants together, regardless of their specific treatment, using a threshold cutoff of 2538.79 µmol·g^−1^ DW to differentiate excluders (below threshold) from non-excluders (above threshold). Both groups included a mix of plants treated with 50 mM and 75 mM NaCl, ensuring that the threshold did not merely classify plants based on treatment levels. The second, the treatment-specific approach, grouped (excluders and non-excluders) by treatment, with threshold values of 1410.44 µmol·g^−1^ DW and 2538.79 µmol·g^−1^ DW for 50 mM and 75 mM treatments, respectively. These thresholds were based on the leaf chloride content of 140 Ru under 50 mM and 75 mM salt treatments, as established in previous studies (Heinitz et al., 2020). In both approaches, the categorization aimed to ensure a robust differentiation between chloride excluders and non-excluders, allowing for an accurate assessment of the predictive capabilities of the PLSDA model.

### 2.5 Stomatal conductance and ϕPS2 measurements

For the second trial, stomatal conductance (g_s_) and the quantum yield of photosystem II (ϕPS2) were measured with a porometer/fluorometer (LI-600, LI-COR Biosciences, Lincoln, NE). One measurement per leaf was recorded between 10 AM and 12 PM on the fifth fully expanded healthy leaf from the base of each plant on the last day of salt treatment. For each measurement, the aperture on the porometer/fluorometer was placed on the right half of the leaf between the veins, and a consistent light intensity (500 ± 50 µmol m⁻² s⁻¹) was maintained on the leaf surface across all samples.

### 2.6 Statistical analysis

A Student’s t-test was used to determine significant differences in chloride content between the two trials. Two-way analysis of variance (ANOVA) was performed to determine treatment, genotype, and their interaction (treatment X genotype) effects. Tukey’s Honest Significant Difference (HSD) was used as a post hoc test to identify significant differences. All the statistical analysis and data visualization were performed using R (version 4.2.1).

## 3 RESULTS

### 3.1 Plant vegetative response to chloride toxicity

Chloride excluder and non-excluder plants growing under 0 mM NaCl treatment did not show any symptoms of salt stress. Under salt stress, different responses were observed. For example, *doaniana* 9026 (*V*. X *doaniana*), a chloride excluder accession, did not show any symptoms and remained green under 75 mM salt treatment (Figure 2A). Conversely, non-excluder accessions, under salt stress (50 mM and 75 mM of NaCl) exhibited wilting and reduced vigor (Figures 2B and 2C). One of the non-excluder accessions, NM11-085 (*V. treleasei*), showed increasing symptoms as stress levels increased; wilting and necrosis were observed at 50 mM salt treatment (Figure 2B), whereas at 75 mM salt treatment, necrosis, wilting, and yellowish leaves were observed (Figure 2C). Generally, brownish-white discoloration started from the margins of the leaves and extended inwards, which is typical of salt stress (Lupo et al., 2023).

**Figure 2.**
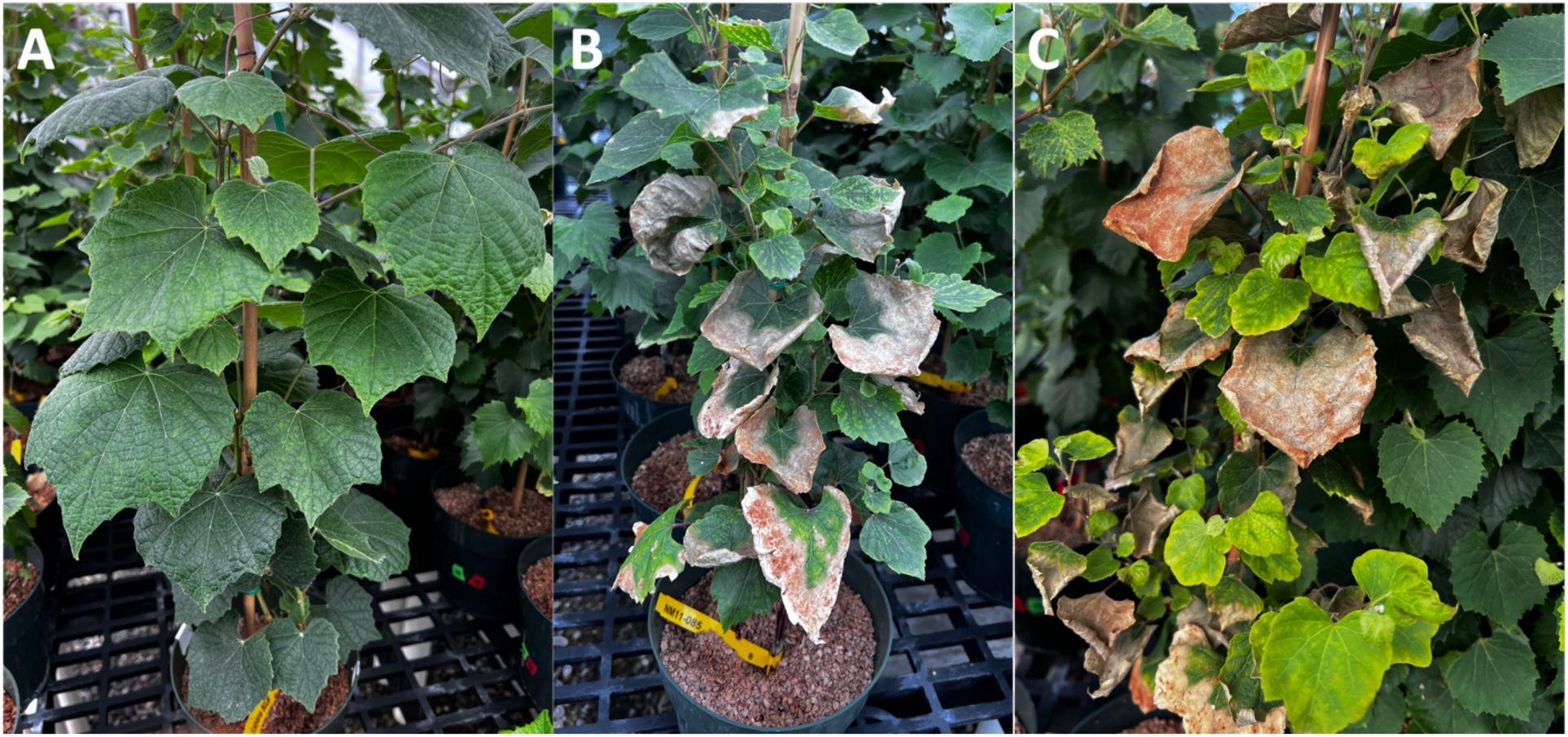
Symptoms of chloride toxicity in chloride-excluding and non-excluding accessions. **(A)** No symptoms of chloride toxicity were observed on the leaves of accession *doaniana* 9026 (*V*. X *doaniana*) under 75 mM salt treatment. Wilting, scorching, and whitish brown discoloration on the leaves of NM11-085 (*V. treleasei*) growing at **(B)** 50 mM and **(C)** 75 mM salt treatments.

### 3.2 Variability in chloride uptake across *Vitis* germplasm

Consistent results were observed across both trials. The chloride content in the first trial ranged from 0 to 7935.97 µmol·g^−1^ DW and from 0 to 7945.27 µmol·g^−1^ DW in the second trial. No significant differences in chloride content between the two trials were observed (*p-value* = 0.17, t-test). There was a high correlation (Pearson’s *r* = 0.93) between trials 1 and 2 for chloride content (Figure 3A).

**Figure 3.**
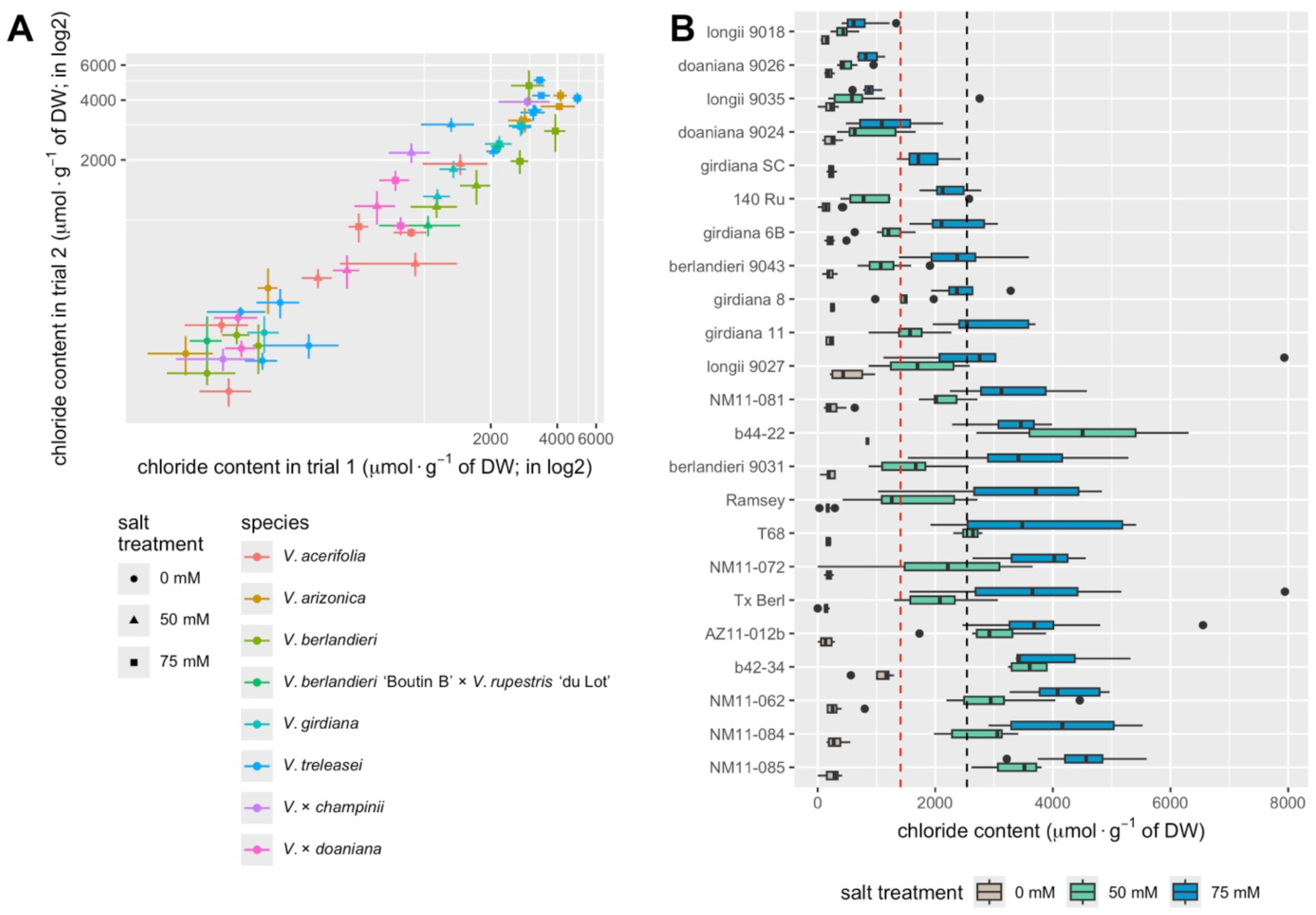
Variability in chloride content accumulation across *Vitis* germplasm. **(A)** Correlation between trials 1 and 2 for leaf chloride; lines represent standard errors for each accession within their corresponding trial. (**B)** Leaf chloride content for 23 accessions under three salt treatments (0, 50, and 75 mM of NaCl). The red dashed line corresponds to the threshold of 1410.44 µmol·g⁻¹ DW, while the black dotted line represents the threshold of 2538.79 µmol·g⁻¹ DW; these thresholds are used to classify excluders and non-excluders in the PLSDA approach.

Significant differences were observed in the treatment, accession, and the interaction between treatment and accession (*p-value <* 0.0001, one-way ANOVA) across both trials. Plants subjected to the 75 mM salt treatment had higher chloride accumulation in their leaves, followed by those treated with 50 mM, and then 0 mM (Figure 3B). longii 9018, a *V. acerifolia* accession, was one of the best chloride excluders found in both trials, with 419.37 µmol·g^−1^ and 713.68 µmol·g^−1^ under the 50 mM and 75 mM treatments, respectively, confirming previous observations (Heinitz et al., 2020). longii 9018 performed comparably to 140Ru (a commercial rootstock derived from *V. rupestris* and *V. berlandieri*), doaniana 9024 (*V.* X *doaniana*), doaniana 9026 (*V.* X *doaniana*), girdiana SC (*V. girdiana*), and longii 9035 (*V. acerifolia*) at 50 mM (Dunnett’s test, *p-value* = 0.05; Figure 3B). However, at 75 mM, the commercial rootstock 140Ru exhibited statistically significantly higher chloride content compared to longii 9018 and the aforementioned chloride excluder accessions, underscoring the importance of other species in acquiring adaptive traits for breeding.

### 3.3 Stomatal conductance and ϕPS2

Stomatal conductance (g_s_) varied between 0.01 to 0.98 mol·m^−2^·s^−1^, with significant differences across varying salt treatments (*p* < 0.001, one-way ANOVA). Plants not subjected to salt stress (0 mM) exhibited the highest g_s_, while no significant differences were detected between the 50 mM and 75 mM treatments (Figure 4A). The ϕPS2 values ranged between 0.06 to 0.75 with significant differences among all treatment groups (*p* < 0.001, one-way ANOVA). The ϕPS2 was significantly higher in plants with no salt treatment (0 mM) and gradually decreased in plants treated with 50 mM and then 75 mM salt (Figure 4B). These observations demonstrated that the salinity treatments applied in these trials induced marked physiological stress, in addition to visible symptoms (Figures 2B and 2C). The correlation (Pearson’s r) between leaf chloride content and both g_s_ and ϕPS2 was −0.41 and −0.40, respectively, illustrating the antagonistic effect of chloride on these physiological traits.

**Figure 4.**
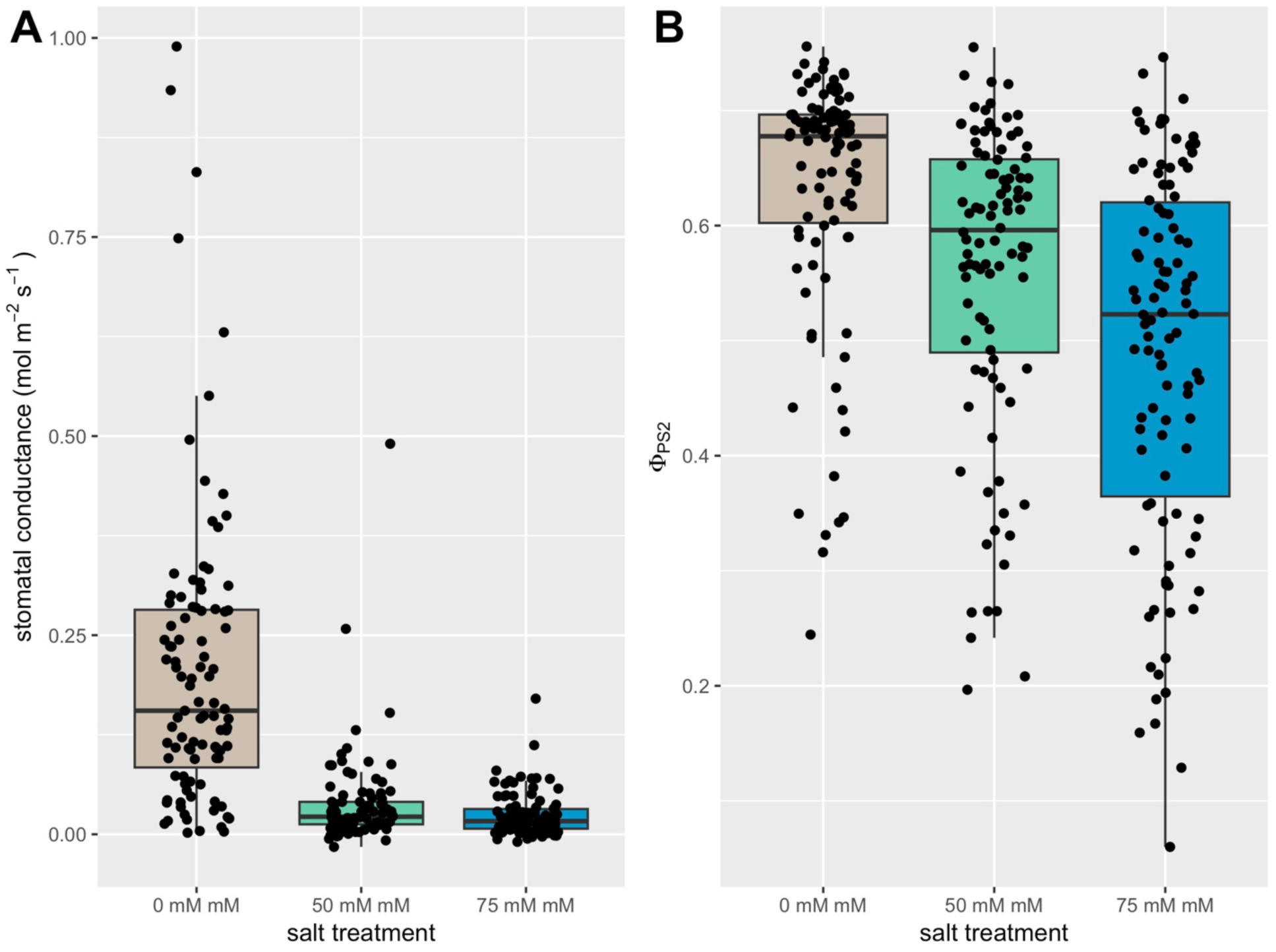
Impact of salt stress on physiological traits. **(A)** stomatal conductance (g_s_), and **(B)** the quantum yield of photosystem II (ϕPS2) under different salt stress treatments; in both panels, points correspond to individual plants.

### 3.4 Exploratory data analysis of spectral data

Variable Importance in Projection (VIP) scores were used to quantify the significance of each wavelength in explaining the variation in the dependent variable, which in this case is chloride content (Figure 5A). Principal component analysis of the spectral data revealed 10 principal components (PCs) explained 99% of the variation. The high amount of variation explained by these PCs is likely due to collinearity among the wavelengths for foliar reflectance.

**Figure 5.**
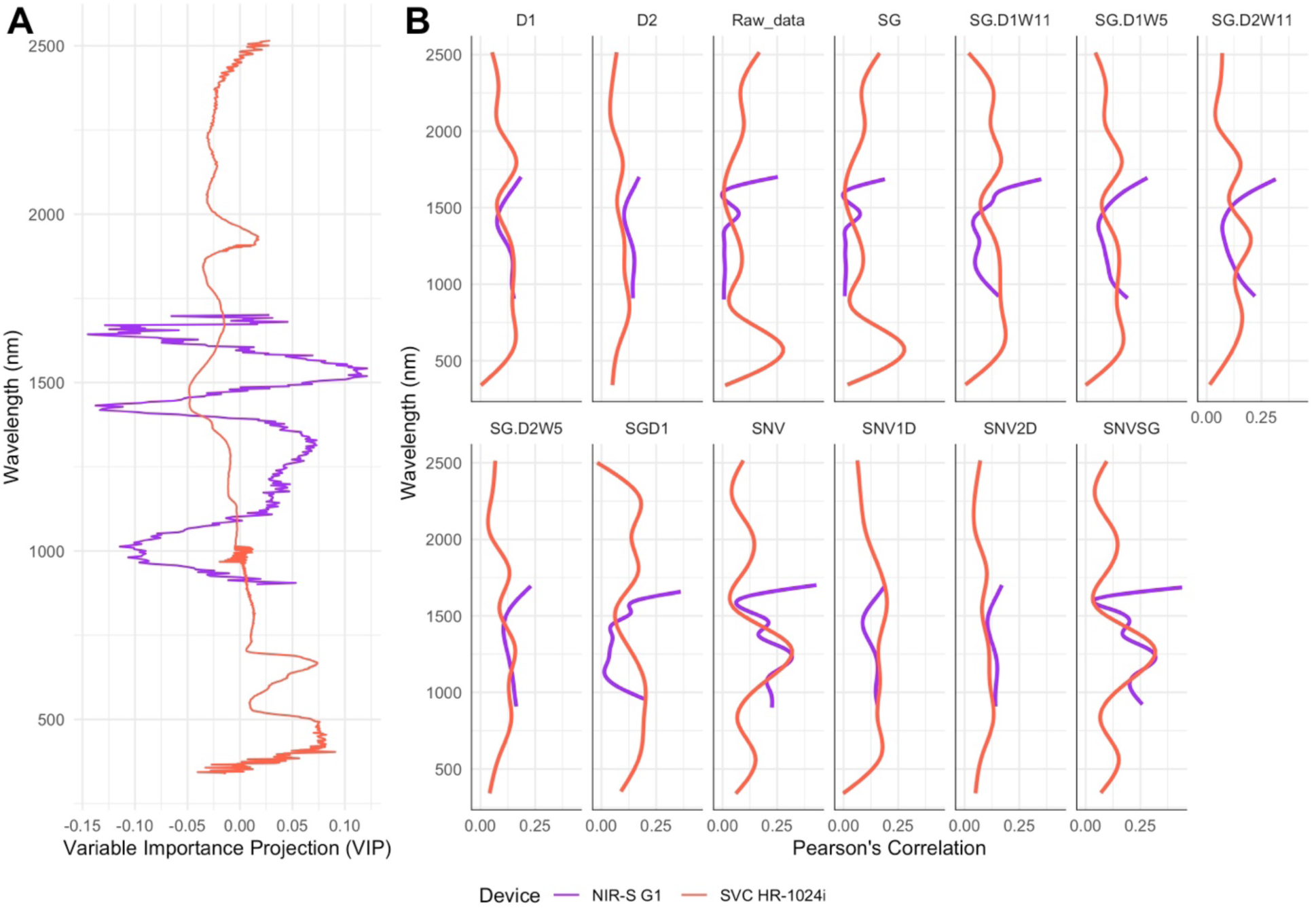
Variable importance in projection (VIP) for chloride content and correlation between chloride content and individual wavelengths. **(A)** VIP estimates for reflectance data obtained with the SVC HR-1024i (red) and NIR-S G1 (purple). **(B)** Pearson’s correlations between chloride content and reflectance at individual wavelengths recorded with the SVC HR-1024i (red) and NIR-S G1 (purple). Pearson’s correlations for different preprocessing along with raw data are shown in different panels. Preprocessing includes first derivative (D1), second derivative (D2), raw data, Savitzky-Golay filter (SG), Savitzky-Golay with window size = 11 and first derivative (SG.D1W11), Savitzky-Golay with window size = 5 and first derivative (SG.D1W5), Savitzky-Golay with window size = 11 and second derivative (SG.D2W11), Savitzky-Golay with window size = 5 and second derivative (SG.D2W5), gap segment derivative with window size = 11 (SGD1), standard normal variate (SNV), standard normal variate and first derivative (SNV1D), standard normal variate and second derivative (SNV2D), and standard normal variate and Savitzky-Golay filter (SNVSG).

The correlations between foliar reflectance at individual wavelengths, as measured by the SVC HR-1024i and the NIR-S G1, and chloride content are detailed in Figure 5B. This includes both raw spectral data and processed spectral data following the 12 preprocessing methods outlined in the Materials and Methods section. For the foliar reflectance from SVC HR-1024i, the strongest correlations between reflectance data and chloride content (Pearson’s *r* = 0.3-0.32) were found within the 613 to 660 nm and 689 to 696 nm ranges, both of which are within the visible light spectrum. Beyond the SNV, SG, and SNVSG, correlations involving other preprocessing varied considerably. Notably, the highest correlation recorded was 0.5 in the 1357-1358 nm region, associated with SNV2D-processed spectral data. Distinct correlation patterns were observed for the data recorded with the NIR-S G1. Specifically, the strongest correlations (Pearson’s *r* = 0.45-0.48) with chloride content were observed in the 1639-1643 nm range for spectra processed with SG.D2W11. Meanwhile, a moderate correlation (Pearson’s *r* = 0.2-0.25) was detected between chloride content and raw spectral data in the 1677-1701 nm region.

### 3.5 Prediction of chloride content using hyperspectral information

Three distinct techniques, PLSR, RF, and SVM, were used to build prediction models to estimate chloride content (as a quantitative variable) using hyperspectral reflectance data. Each technique was tested using the raw reflectance data as well as the spectra processed with the 12 different preprocessing discussed above, resulting in 39 different models (13 preprocessing x 3 techniques). Several metrics, including *R^2^*, *RMSE, RPIQ, CCC,* and bias were used to evaluate the prediction models. The best model was selected based on the highest *R^2^*, lowest *RMSE*, and bias closest to zero.

Five different scenarios were analyzed, which included both model training and validation within individual trials, combined trials, and across trials (Figure 1). These analyses utilized only the spectral data from the SVC HR-1024i device, as it was the only dataset recorded in both trials. For data recorded with the NIR-S G1, only scenario 1 was tested. Based on *R^2^*, *RMSE,* and bias values, spectral data processed with SNVSG consistently produced the most accurate prediction models across all evaluated scenarios (Figure 6A, TableS2). When the SNVSG was applied, the PLSR models outperformed the others, showing the highest average *R^2^* (0.67), followed by SVM models (0.53), and RF models (0.36).

**Figure 6.**
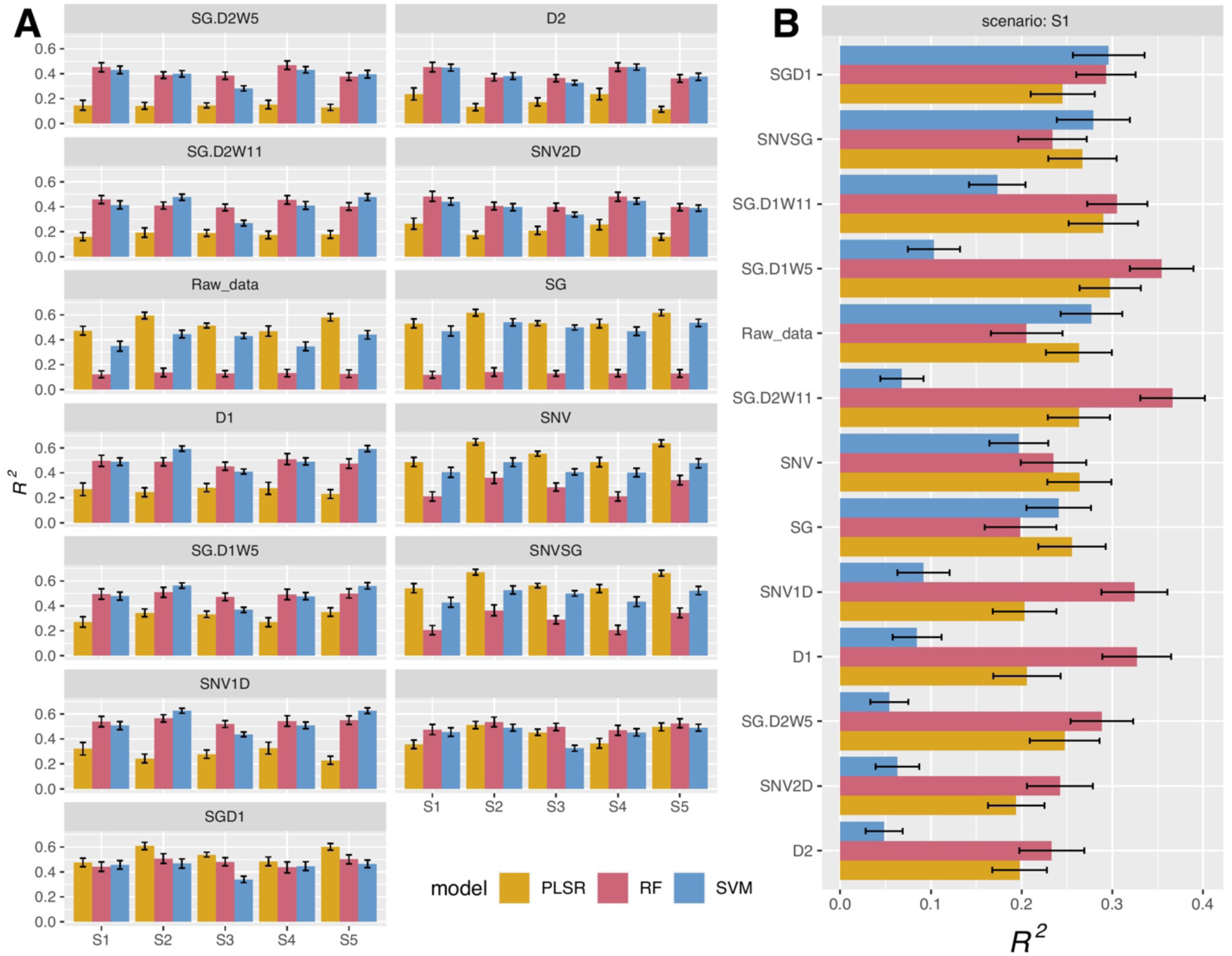
Comparison of modeling approaches for predicting chloride content across different training and validation scenarios. **(A)** 12 preprocessing methods along with raw data obtained using SVC HR-1024i shown in each subpanel with different scenarios shown on the X-axis and squared Pearson’s correlation between observed and predicted values in the validation set (*R^2^*) on the Y-axis. **(B)** Raw data along with 12 preprocessing methods applied to spectral data from NIR-S G1 shown in the Y-axis and squared Pearson’s correlation between observed and predicted values in the validation set (*R^2^*) displayed in the X-axis.

In scenario 1, the PLSR model employing SNVSG processing attained an *R^2^* of 0.54 and an *RMSE* of 697 µmol·g^−1^ DW. Among the 12 preprocessing methods applied to the reflectance data, the PLSR model performed best in five, the RF model in eight, and the SVM model in none of the preprocessing methods applied. Despite this, the PLSR models surpassed both the RF and SVM models in performance, evidenced by achieving the highest *R^2^* and the lowest *RMSE* values. For scenario 2, the PLSR model with SNVSG preprocessing marked the highest *R^2^* at 0.67 and the lowest *RMSE* at 574.1 µmol·g^−1^ DW. Out of the twelve preprocessing methods, the PLSR model was superior in five, the SVM in six, and the RF in two preprocessing methods. In scenario 3, the PLSR model again led with an *R^2^* of 0.56 and an *RMSE* of 665.2 µmol·g^−1^ DW with spectra processed with SNVSG. Here, the PLSR model showed superior performance in five, the RF in eight, and the SVM in none of the preprocessing methods applied. In scenario 4, the PLSR model reached the highest *R^2^* of 0.54 and the lowest *RMSE* of 691.83 µmol·g^−1^ DW with SNVSG. The PLSR model was superior in five, RF in eight, and SVM in none of the preprocessing methods applied. Lastly, in scenario 5, the PLSR model achieved a peak mean *R^2^* of 0.66 and a mean *RMSE* of 580.2 µmol·g^−1^ DW with SNVSG. The performance rankings among the preprocessing methods were similar to those in scenario 2, with the PLSR model leading in five, the SVM in six, and the RF in two. The details on *R^2^* values, *RMSE, RPD, RPIQ, CCC,* and bias for three techniques and 12 preprocessing for five scenarios are presented in Table S2.

While the SNVSG yielded the highest prediction accuracies for data recorded with the SVC HR-1040i using PLSR, the best predictions for the NIR-S G1 (Figure 6B) were achieved with RF with SG.D2W11 (*R^2^* = 0.37, *RMSE* = 1236.82 µmol·g^−1^ DW), followed by SG.D1W5 (*R^2^* = 0.35, *RMSE* = 1245.58 µmol·g^−1^ DW), and D1 (*R^2^* = 0.33, *RMSE* = 1276.08 µmol·g^−1^ DW).

### 3.6 Hyperspectral predictions with a classification model

While we showed that hyperspectral data can be used to predict chloride content (Figures 5B, 6A, and 6B), the accuracy of these predictions was not yet optimal. Therefore, a classification-based approach was explored using PLSDA. For this analysis, chloride content from plants grown at 50 and 75 mM was categorized into two groups: chloride excluders (leaf chloride content below threshold), and non-excluders (leaf chloride content above threshold). The threshold was based on chloride content for 140Ru under 75 mM (2538.79 µmol·g^−1^ DW). This classification resulted in 216 plants being identified as excluders (132 plants from treatment 50 mM and 84 plants from treatment 75 mM), and 158 as non-excluders (51 plants treated with 50 mM and 107 with 75 mM). The only scenario tested was scenario 3, where 70% of the data recorded in both trials was used for model training and the remaining 30% for validation. For the SVC HR-1024i, the average accuracy ranged from 0.83 to 0.89, depending on the preprocessing method applied, whereas specificity ranged from 0.85 to 0.90 (Figure 7, Table S3). With spectral data from the NIR-S G1, average accuracy varied from 0.69 to 0.97, and sensitivity ranged from 0.77 to 0.98 (Figure 7).

**Figure 7.**
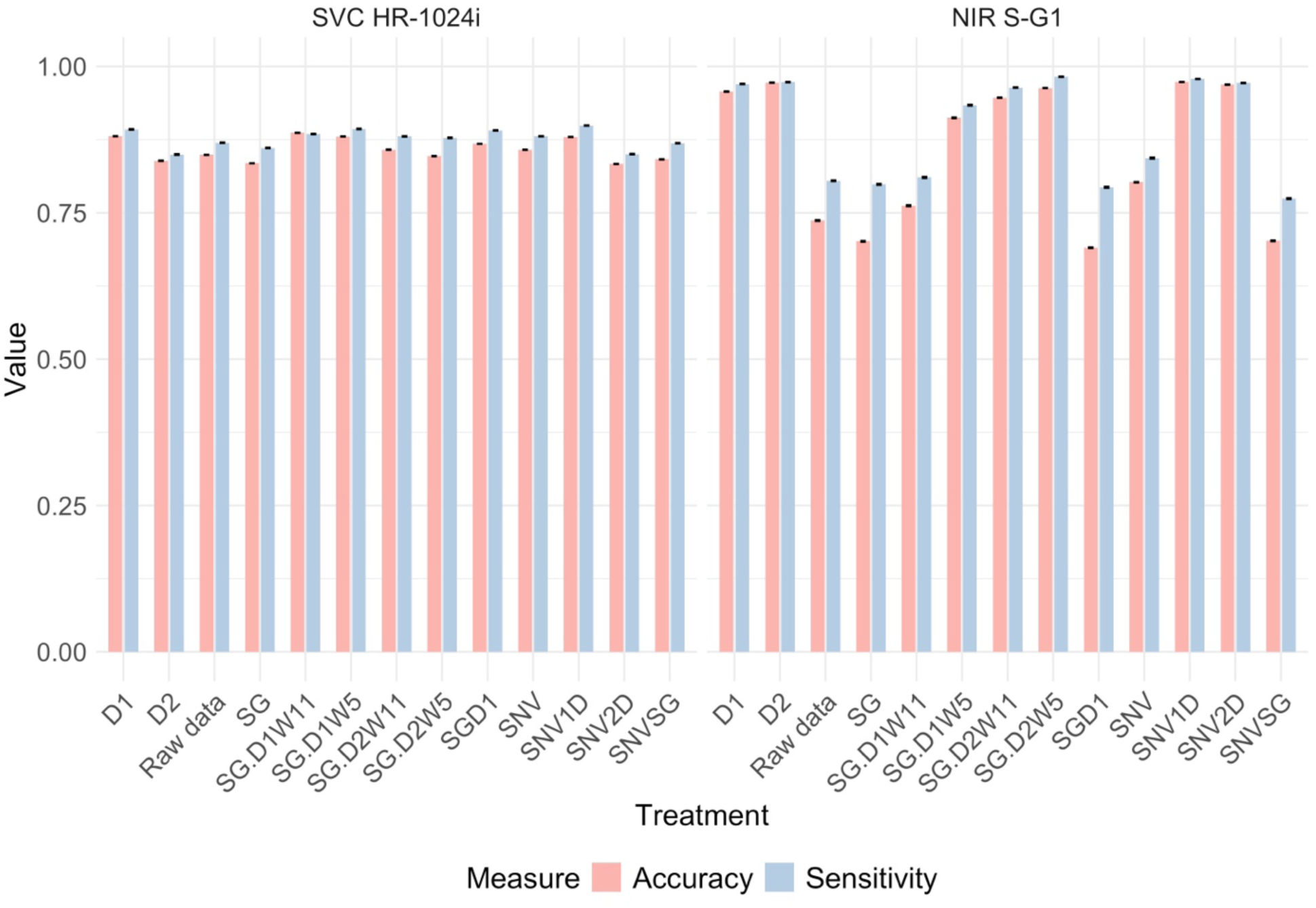
Accuracy and specificity results for PLSDA with raw data and 12 preprocessing methods for both devices - SVC HR-1024i and NIR-S G1.

A similar analysis was conducted for each salt treatment separately; that is, excluders and non-excluders were identified within each treatment and used to build prediction models. The thresholds for discretizing within each treatment were 1410.44 µmol·g^−1^ and 2538.79 µmol·g^−1^ DW for the 50 mM and 75 mM treatments, respectively. For the 50 mM treatment, 104 plants were categorized as non-excluders, and 79 as excluders. Similarly, for the 75 mM treatment, 107 plants were classified as non-excluders and 84 as excluders. Accuracies ranged from 0.91 to 0.99 and 0.74 to 1 and sensitivity ranged from 0.89 to 0.99 and 0.73 to 1 for SVC HR-1024i and NIR-S G1, respectively, across the 12 preprocessing methods tested (Tables S4 and S5).

## 4 Discussion

### 4.1 Impacts of salinity stress on leaves

Increasing NaCl concentration in the treatments led to higher chloride content in the leaves. Genotypes treated with no salt (0 mM) showed no variation in chloride content, regardless of their classification as excluders or non-excluders (Figure 4B). Chloride excluders, such as longii 9018, treated with 50 mM and 75 mM NaCl accumulated similar chloride content as under 0 mM for the same accession. However, non-excluders accumulated significantly higher chloride content in their leaves under 50 mM and 75 mM NaCl treatments compared to 0 mM.

Chloride tolerance in *Vitis* is not fully understood, as various mechanisms are believed to exist across different species. Some species prevent chloride uptake in the leaves, instead of accumulating it in the root system, while others show no chloride accumulation in either plant organ (Heinitz et al., 2020; Cochetel et al., 2023). For instance, the chloride-excluder accession 140 Ru has been observed to accumulate less chloride in the leaves and more in the roots, due to decreased xylem loading and restricted root-to-shoot chloride transport (Tregeagle et al., 2010; Henderson et al., 2014; Heinitz et al., 2020). Conversely, *V. girdiana*, another chloride excluder, was found to have low chloride content in both leaves and roots (Heinitz et al., 2020).

Water and salt stress induce osmotic stress, leading to stomatal closure and subsequent reduction in gas exchange (Kusvuran, 2012; Baneh et al., 2014; Liao et al., 2022). Therefore, measuring stomatal conductance can serve as an indicator of stress in plants. In our experiment, the stomatal conductance in plants under salt treatment at 50 and 75 mM of NaCl was significantly lower than in those under no salt (0 mM). No differences in stomatal conductance were observed for both stress treatments, suggesting that 50 mM of salt was sufficient to promote stomatal closing and reduced gas exchange. There were no significant differences observed between excluders and non-excluders for stomatal conductance and ϕPS2 under 50 and 75 mM, highlighting that the osmotic phase rather than the ionic phase of the salt stress led to the stomatal closure. However, a stepwise decrease was observed in ϕPS2 at each treatment than in stomatal conductance, which could be related to the decoupling of the light and carbon reactions of photosynthesis (Magney et al., 2020). Similar results have been documented in other grape cultivars, such as ‘Soltanin’ and ‘Fakhri’ when salinity increased from 0 to 250 mM (Bybordi, 2012). Furthermore, chlorophyll a and b content decreased as the salinity increased with the lowest values observed at 250 mM (Bybordi, 2012). Comparable reductions have also been documented in four native species (Paudel and Sun, 2023) and Ginkgo biloba (Zhao et al., 2019). Given the correlations between salt stress and both ϕPS2 and stomatal conductance, these parameters can be utilized as proxies to assess plant stress and the resultant impact on photosynthetic machinery and gas exchange (Maxwell and Johnson, 2000).

### 4.2 Comparison between different models and preprocessing methods

Raw reflectance data are often noisy and require further processing to increase the signal-to-noise ratio, reduce variation introduced by light scattering, and improve resolution. The 12 spectral preprocessing methods used in this study highlighted different features of the spectral profiles, impacting the accuracy of the predictions. For instance, SNV corrects for light scattering and other multiplicative interferences such as particle size variations in the sample (Grisanti et al., 2018). SG filtering preserves important features like the width and height of spectral peaks while minimizing the influence of random noise, making it useful for analyzing noisy spectrometric data (Delwiche et al., 2010). Derivatives (either first derivative, D1, or second derivative, D2) can highlight subtle features and significant variations in the spectral curves (Rinnan et al., 2009; Dotto et al., 2018). D1 reduces background trends that do not have sharp changes, making spectral peaks more distinct, whereas D2 makes it easier to locate inflection points where the curvature of the spectrum changes.

Among the three machine learning models built with the SVC HR-1024i data, PLSR had the highest *R^2^* of 0.67. PLSR consistently produced higher *R^2^* values compared to RF and SVM when using raw data, SNV, SG, or a combination of SNV and SG for data processing. Although RF and SVM performed better than PLSR when derivatives were used for data processing, the highest *R^2^* value was with the PLSR model shown in Figure 6A. The *R^2^* values with derivatives were comparatively lower than SNV or SG, highlighting that the PLSR model is better at handling spectral data overall. PLSR is effective with both wide data (many variables with few samples) and tall data (fewer variables with a larger number of samples), and it excels in managing highly correlated spectral data. In contrast, SVM is better suited for modeling non-linear relationships (Thissen et al., 2004). RF is also better when handling non-linear relationships and larger data sets. PLSR models using the SNVSG transformation consistently outperformed other models by minimizing noise and light scattering (Figure 6A). Notably, the model trained on trial 2 and validated on trial 1 yielded an *R^2^* value of 0.66 and the lowest *RMSE* value of 891.11 µmol·g^−1^ DW, indicating effective applicability across trials. The higher *R^2^* values observed under scenario 2 can likely be attributed to the screening of a slightly larger number of plants in trial 2, as 38 plants did not survive in trial 1. Additionally, training the model with data from trial 2, which included more plants, and testing it on a slightly smaller dataset from trial 1 likely contributed to the enhanced *R^2^* observed in scenario 5. With the use of spectral data from NIR-S G1, the RF model using the SG.D2W11 performed better with an *R^2^* value of 0.37 (Figure 6B). Derivative preprocessing likely performed better because it highlighted subtle features, revealing small but significant differences in the spectral data. The variation in model performance with different devices may be attributed to factors such as the sensor, slit size, light source, hardware, and other features specific to each device. Therefore, careful consideration should be given to identifying the appropriate preprocessing method based on the spectral data, device type, and the trait of interest (Xiao et al., 2022; Nkouaya Mbanjo et al., 2022; Hershberger et al., 2022).

Compared to the NIR-S-G1, the SVC HR-1024i demonstrated greater sensitivity to pigments such as chlorophyll, carotenoids, and anthocyanins within the visible spectrum (VIS), which interact with chloride in the leaves (Blackburn, 2007). In our analysis, the strongest correlations (Pearson’s *r*) between chloride content and wavelengths were found within the 613-660 nm and 689-696 nm range (for raw spectral data recorded with the SVC HR-1024i). In a similar study conducted in persimmon trees, the 390-472 nm and 690-692 nm wavelength ranges were also strongly associated with chloride accumulation in leaves (Visconti and de Paz, 2019). Chloride ions in the leaves are known to interact with pigments like chlorophyll a, chlorophyll b, and β carotene, which affect how light is absorbed, and in turn, reflected. The reduced ϕPS2 in plants under salinity stress suggests altered pigment composition and/or changes in light energy balance (Maxwell and Johnson, 2000). While improvements in hyperspectral-based predictions are possible, as demonstrated by the study in persimmon conducted by Visconti and De Paz (2019), which utilized a benchtop spectrometer and dried grounded samples to achieve an *R^2^* value of 0.93, the ability to screen and process a larger quantity of plants—even with slight reductions in accuracy— may be preferred in certain applications such as plant breeding.

The analysis of spectral data with PLSDA, aiming to predict classes (chloride excluder vs. non-excluder), provided consistently better accuracy and sensitivity results with the SVC HR-1024i regardless of the preprocessing method. The use of NIR-S G1 spectral data for PLSDA outperformed the SVC HR-1024i when derivatives were applied. This might be because derivatives enhanced subtle but significant differences in reflectance between excluders and non-excluders. Misclassification with PLSDA may arise due to an overlap in chloride content between excluders and non-excluders. Additionally, chloride content is highly influenced by environmental factors, causing threshold values to vary across different years and environments. Therefore, future research should focus on screening more accessions and environments to optimize PLSDA for classification.

### 4.3 Application of Hyperspectral Sensing to Grape Breeding Programs

The capability to screen thousands of plants during the breeding process can increase genetic gain by allowing for higher selection intensities. However, given the typically constrained budgets of breeding programs, breeders must consider the cost-effectiveness of new screening methodologies relative to the expected improvement in genetic gain. SVC HR-1024i performs better for predicting the values whereas NIR-S G1 achieves comparably high accuracy when the objective is simplified to a classification problem (chloride excluder vs. non-excluder), despite its relatively low price. In practical terms, a grapevine breeding program could significantly increase the volume of plants screened for chloride exclusion while keeping screening costs low, given that the investment in equipment remains minimal.

## AUTHOR CONTRIBUTIONS

Sadikshya Sharma: Conceptualization; methodology; data collection; formal analysis; visualization; data acquisition; writing—original draft. Christopher Wong: data acquisition; formal analysis; visualization; writing – review and editing. Krishna Bhattarai: data collection, formal analysis; visualization; writing – review, and editing. Yaniv Lupo: methodology; formal analysis; visualization. Troy Magney: data acquisition; formal analysis; visualization; writing – review and editing. Luis Diaz-Garcia: project administration; funding acquisition; formal analysis; writing—review and editing

## ACKNOWLEDGMENTS

The authors would like to thank Veronica Nunez, Christopher Chen, Claire Heinitz, Sabrina Colacion, and Mikayla Bailey for their support during propagation and greenhouse management.

## DATA AVAILABILITY STATEMENT

All the collected data have been made available in the supplementary files accompanying this manuscript.

## References

Asner, G. P. (1998). Biophysical and biochemical sources of variability in canopy reflectance. Remote sensing of Environment, 64(3), 234–253. 10.1016/S0034-4257(98)00014-5

Baneh, H. D., Attari, H., Hassani, A., Abdollahi, R., Taheri, M., & Ghani, F. (2014). Genotypic variation in plant growth and physiological response to salt stress in grapevine. Philipp. Agr. Sci, 97(2), 113–121.

Blackburn, G. A. (2007). Hyperspectral remote sensing of plant pigments. Journal of experimental botany, 58(4), 855–867.

Bybordi, A. (2012). Study effect of salinity on some physiologic and morphologic properties of two grape cultivars. Life Science Journal, 9(4), 1092–1101.

Calzone, A., Cotrozzi, L., Lorenzini, G., Nali, C., & Pellegrini, E. (2021). Hyperspectral detection and monitoring of salt stress in pomegranate cultivars. Agronomy, 11(6), 1038. 10.3390/agronomy11061038

Cochetel, N., Minio, A., Guarracino, A., Garcia, J.F., Figueroa-Balderas, R., Massonnet, M., Kasuga, T., Londo, J.P., Garrison, E., Gaut, B.S. & Cantu, D. (2023). A super-pangenome of the North American wild grape species. Genome Biology, 24(1), 290. 10.1186/s13059-023-03133-2

Delwiche, S. R., & Reeves, J. B. (2010). A graphical method to evaluate spectral preprocessing in multivariate regression calibrations: Example with Savitzky–Golay filters and partial least squares regression. Applied spectroscopy, 64(1), 73–82.

de Paz, J. M., Visconti, F., Chiaravalle, M., & Quiñones, A. (2016). Determination of persimmon leaf chloride contents using near-infrared spectroscopy (NIRS). Analytical and Bioanalytical Chemistry, 408, 3537–3545. 10.1007/s00216-016-9430-2

Dotto, A. C., Dalmolin, R. S. D., ten Caten, A., & Grunwald, S. (2018). A systematic study on the application of scatter-corrective and spectral-derivative preprocessing for multivariate prediction of soil organic carbon by Vis-NIR spectra. Geoderma, 314, 262–274. 10.1016/j.geoderma.2017.11.006

Gates, D. M., Keegan, H. J., Schleter, J. C., & Weidner, V. R. (1965). Spectral properties of plants. Applied optics, 4(1), 11–20.

Grisanti, E., Totska, M., Huber, S., Krick Calderon, C., Hohmann, M., Lingenfelser, D., & Otto, M. (2018). Dynamic localized SNV, Peak SNV, and partial peak SNV: Novel standardization methods for preprocessing of spectroscopic data used in predictive modeling. Journal of Spectroscopy, 2018(1), 5037572. 10.1155/2018/5037572

Heinitz, C. C., Riaz, S., Tenscher, A. C., Romero, N., & Walker, M. A. (2020). Survey of chloride exclusion in grape germplasm from the southwestern United States and Mexico. Crop Science, 60(4), 1946–1956. 10.1002/csc2.20085

Henderson, S. W., Baumann, U., Blackmore, D. H., Walker, A. R., Walker, R. R., & Gilliham, M. (2014). Shoot chloride exclusion and salt tolerance in grapevine is associated with differential ion transporter expression in roots. BMC plant biology, 14, 1–18. 10.1186/s12870-014-0273-8

Hershberger, J., Mbanjo, E.G.N., Peteti, P., Ikpan, A., Ogunpaimo, K., Nafiu, K., Rabbi, I.Y. & Gore, M.A. (2022). Low-cost, handheld near-infrared spectroscopy for root dry matter content prediction in cassava. The Plant Phenome Journal, 5(1), e20040. 10.1002/ppj2.20040

Hershberger, J., Morales, N., Simoes, C. C., Ellerbrock, B., Bauchet, G., Mueller, L. A., & Gore, M. A. (2021). Making waves in Breedbase: An integrated spectral data storage and analysis pipeline for plant breeding programs. The Plant Phenome Journal, 4(1), e20012. 10.1002/ppj2.20012

Huang, J., Liao, H., Zhu, Y., Sun, J., Sun, Q., & Liu, X. (2012). Hyperspectral detection of rice damaged by rice leaf folder (Cnaphalocrocis medinalis). Computers and electronics in agriculture, 82, 100–107. 10.1016/j.compag.2012.01.002

Jin, X., Chen, X., Shi, C., Li, M., Guan, Y., Yu, C.Y., Yamada, T., Sacks, E.J. & Peng, J., 2017 (2017). Determination of hemicellulose, cellulose and lignin content using visible and near infrared spectroscopy in Miscanthus sinensis. Bioresource technology, 241, 603–609. 10.1016/j.biortech.2017.05.047

Knipling, E. B. (1970). Physical and physiological basis for the reflectance of visible and near-infrared radiation from vegetation. Remote sensing of environment, 1(3), 155–159.

Kokaly, R. F., & Skidmore, A. K. (2015). Plant phenolics and absorption features in vegetation reflectance spectra near 1.66 μm. International Journal of Applied Earth Observation and Geoinformation, 43, 55–83. 10.1016/j.jag.2015.01.010

Kusvuran, S. (2012). Effects of drought and salt stresses on growth, stomatal conductance, leaf water, and osmotic potentials of melon genotypes (Cucumis melo L.). African Journal of Agricultural Research, 7(5), 775–781.

Liao, Q., Gu, S., Kang, S., Du, T., Tong, L., Wood, J. D., & Ding, R. (2022). Mild water and salt stress improve water use efficiency by decreasing stomatal conductance via osmotic adjustment in field maize. Science of the Total Environment, 805, 150364. 10.1016/j.scitotenv.2021.150364

Lupo, Y., Prashanth, K., Lazarovitch, N., Fait, A., & Rachmilevitch, S. (2023). Importance of leaf age in grapevines (Vitis spp.) under salt stress. Scientia Horticulturae, 321, 112325. 10.1016/j.scienta.2023.112325

Magney, T. S., Barnes, M. L., & Yang, X. (2020). On the covariation of chlorophyll fluorescence and photosynthesis across scales. Geophysical Research Letters, 47(23), e2020GL091098. 10.1029/2020GL091098

Maxwell, K., & Johnson, G. N. (2000). Chlorophyll fluorescence—a practical guide. Journal of experimental botany, 51(345), 659–668.

Niederberger, J., Todt, B., Boča, A., Nitschke, R., Kohler, M., Kühn, P., & Bauhus, J. (2015). Use of near-infrared spectroscopy to assess phosphorus fractions of different plant availability in forest soils. Biogeosciences, 12(11), 3415–3428. 10.5194/bg-12-3415-2015

Nkouaya Mbanjo, E.G., Hershberger, J., Peteti, P., Agbona, A., Ikpan, A., Ogunpaimo, K., Kayondo, S.I., Abioye, R.S., Nafiu, K., Alamu, E.O. & Adesokan, M. (2022). Predicting starch content in cassava fresh roots using near-infrared spectroscopy. Frontiers in Plant Science, 13, 990250. 10.3389/fpls.2022.990250

Olesen, J.E., Trnka, M., Kersebaum, K.C., Skjelvåg, A.O., Seguin, B., Peltonen-Sainio, P., Rossi, F., Kozyra, J. & Micale, F. (2011). Impacts and adaptation of European crop production systems to climate change. European journal of agronomy, 34(2), 96–112. 10.1016/j.eja.2010.11.003

Ollat, N., Peccoux, A., Papura, D. & Esmenjaud, D., (2016). Rootstocks as a component of adaptation to environment. In ‘Grapevine in a changing environment: a molecular and ecophysiological perspective’. (Eds H Gerós, M Manuela, C Hipólito, M Gil, S Delrot) pp. 68–108.

Paudel, A., & Sun, Y. (2023). Growth, Morphological, and Biochemical Responses of Four Native Species to Salinity Stress. HortScience, 58(6), 651–659. 10.21273/HORTSCI17044-23

Rinnan, Å., Van Den Berg, F., & Engelsen, S. B. (2009). Review of the most common pre-processing techniques for near-infrared spectra. TrAC Trends in Analytical Chemistry, 28(10), 1201–1222. 10.1016/j.trac.2009.07.007

Thissen, U., Pepers, M., Üstün, B., Melssen, W. J., & Buydens, L. M. C. (2004). Comparing support vector machines to PLS for spectral regression applications. Chemometrics and Intelligent Laboratory Systems, 73(2), 169–179. 10.1016/j.chemolab.2004.01.002

Tregeagle, J. M., Tisdall, J. M., Tester, M., & Walker, R. R. (2010). Cl–uptake, transport and accumulation in grapevine rootstocks of differing capacity for Cl–-exclusion. Functional Plant Biology, 37(7), 665–673. 10.1071/FP09300

Visconti, F., & de Paz, J. M. (2019). Non-destructive assessment of chloride in persimmon leaves using a miniature visible near-infrared spectrometer. Computers and Electronics in Agriculture, 164, 104894. 10.1016/j.compag.2019.104894

Walker, R. R. (1994). Grapevine responses to salinity. TT-Reaction de la vigne a la salinite. Bulletin de OIV, 76, 634–661.

Walker, R. R., Blackmore, D. H., Clingeleffer, P. R., Godden, P., Francis, L., Valente, P., & Robinson, E. (2003). Salinity effects on vines and wines.

Wong, C. Y. (2023). Plant optics: underlying mechanisms in remotely sensed signals for phenotyping applications. AoB Plants, 15(4), plad039. 10.1093/aobpla/plad039

Wong, C. Y., Gilbert, M. E., Pierce, M. A., Parker, T. A., Palkovic, A., Gepts, P., … & Buckley, T. N. (2023). Hyperspectral remote sensing for phenotyping the physiological drought response of common and tepary bean. Plant Phenomics, 5, 0021. DOI: 10.34133/plantphenomics.002

Xiao, Q., Tang, W., Zhang, C., Zhou, L., Feng, L., Shen, J., Yan, T., Gao, P., He, Y. & Wu, N., 2022 (2022). Spectral preprocessing combined with deep transfer learning to evaluate chlorophyll content in cotton leaves. Plant Phenomics. DOI: 10.34133/2022/981384

Zhang, J., Zhang, W., Xiong, S., Song, Z., Tian, W., Shi, L., & Ma, X. (2021). Comparison of new hyperspectral index and machine learning models for prediction of winter wheat leaf water content. Plant Methods, 17(1), 1–14. 10.1186/s13007-021-00737-2

Zhao, H., Liang, H., Chu, Y., Sun, C., Wei, N., Yang, M., & Zheng, C. (2019). Effects of salt stress on chlorophyll fluorescence and the antioxidant system in Ginkgo biloba L. seedlings. HortScience, 54(12), 2125–2133. 10.21273/HORTSCI14432-19

